# Identification of individual differences in response to methadone, buprenorphine, and naltrexone in animal models of opioid use disorder

**DOI:** 10.1101/2022.07.23.501217

**Authors:** Marsida Kallupi, Giordano de Guglielmo, Dana Conlisk, Molly Brennan, Lani Tieu, Sharona Sedighim, Brent Boomhower, Lauren C Smith, Kokila Shankar, Lieselot LG Carrette, Sierra Simpson, Alicia Avelar, Lisa Maturin, Angelica Martinez, Ran Qiao, Selen Dirik, Caitlin Crook, Selene Bonnet-Zahedi, Mohini R. Iyer, Corrine E. Blucher, McKenzie J Fannon, Leah C. Solberg Woods, Abraham A. Palmer, Olivier George

**Affiliations:** Department of Psychiatry, University of California, San Diego, La Jolla CA 92093, USA; Department of Neuroscience, The Scripps Research Institute, La Jolla, CA 92037, USA; Wake Forest University Health Sciences, Winston-Salem, NC 27157, US; Institute for Genomic Medicine, University of California, San Diego, La Jolla CA 92093, USA

**Keywords:** methadone, buprenorphine, naltrexone, oxycodone, escalation, hyperalgesia, analgesia, motivation

## Abstract

**Rationale:** Current medications for opioid use disorder include buprenorphine, methadone, and naltrexone. While these medications show significant efficacy in reducing craving and opioid use, there are substantial individual differences in response to these treatments in humans. The reason for such difference is poorly known.

**Objectives:** Here, we tested the hypothesis that similar individual differences may be observed in a large population of heterogenous stock rats, that have been bred to maximize genetic diversity, using a behavioral paradigm relevant to opioid use disorder.

**Methods:** Over 500 rats were given intermittent (4d/week) and extended access (12h/day) to oxycodone self-administration for 14 sessions to establish oxycodone dependence and escalation of intake. We then measured the effect of buprenorphine (0.5mg/kg), methadone (3mg/kg) and naltrexone (3mg/kg) on the motivation to self-administer oxycodone by using a progressive ratio schedule of reinforcement.

**Results:** We found that naltrexone and buprenorphine significantly decreased motivation to oxycodone rewards. While naltrexone reduced oxycodone intake in both males and females, systemic administration with buprenorphine reduced progressive ratio responses only in males. Methadone reduced motivation to oxycodone self-administration in nearly 25% of the population, without reaching statical significance. Our results showed that the efficacy of these medications depends on the severity of addiction like behaviors, indicated by the addiction index.

**Conclusions:** These results demonstrate individual differences in response to medications to treat opioid use disorder in a genetically diverse population of rats.

## Introduction

Although opioid medications effectively treat acute pain and help relieve chronic pain for some patients (Moore et al., 2013), their use presents a dilemma for healthcare providers because of the risk of addiction. In the last decade, the opioid epidemic has become a pervasive crisis, with negative effects not only on the individuals but on the entire society. Many efforts have focused on identifying mechanisms that underlie vulnerability to opioid abuse.

One of the most used analgesic compounds available for treating moderate-to-severe pain is oxycodone. While oxycodone appears to be equally effective to morphine in some types of pain (e.g., post-operative pain and cancer pain) [1, 2], clinical data demonstrate preference for oxycodone consumption in comparison with other medical and non-medical opioids [3].

To date, three treatments have been approved by the United States Food and Drug Administration for the treatment of opioid use disorder (OUD): methadone, buprenorphine, naltrexone. Methadone is a synthetic opioid compound that binds primarily to the MOP-opioid receptor (MOP). This pharmacological profile of methadone raises concerns and limits its use, as it can cause a dose-dependent decrease in respiration often associated with apnea at high doses. Buprenorphine is a long-lasting partial agonist at MOP and NOP receptors, and an antagonist at DOP and KOP receptors [4, 5]. Methadone and buprenorphine are particularly effective in reducing opioid-induced mortality and maintaining patients in treatment, but important safety concerns and strict regulations because of their agonist properties at MOPs have limited their use [6, 7]. Substitution therapy with buprenorphine or methadone reduces both somatic and affective signs of withdrawal by maintaining opioid dependence. Therefore, a better understanding of the neurobiological mechanisms of environmental stimuli and internal states associated with opioid withdrawal and the alleviation of withdrawal may help identify previously unidentified strategies to decrease relapse propensity in individuals who have been abstinent for weeks, months, or even years. Naltrexone is a long-lasting opioid antagonist, and unlike buprenorphine or methadone, it is not associated with the development of tolerance and dependence, and lacks the potential for misuse [8, 9]. The pharmacological properties of naltrexone limit however its effectiveness. The relatively short duration of action and the competitive binding to the receptor, may be compromised by administration of large doses of an agonist and often lead to precipitated withdrawal. In summary, while the current medications are effective for opioid abuse in some individuals, there is still a significant need for better approaches to treat opioid addiction [10-13].

Several studies have examined the effects of FDA approved drugs on the rewarding potency, extinction, and relapse vulnerability from opioids such as oxycodone, heroine or fentanyl [5, 14-19].

Preclinical and clinical evidence suggests that opioids need individual dose titration, and each individual presents different profiles of tolerability [20]. In this study, we used heterogenous stock rats (HS) and a preclinical model of oxycodone dependence to investigate the effect of acute buprenorphine, methadone and naltrexone treatment on the motivation to self-administer oxycodone in a progressive ratio test, in male and female HS rats. HS rats are a unique outbred strain of rats, characterized by high genetic variability that has been developed to mimic genetic variability in humans. HS rats are the most highly recombinant rat intercross available and are a powerful tool for genetic studies [21, 22]. HS rats were initially trained to self-administer oxycodone (0.5 ug/ml) daily in ShA schedule (2h), then moved to LgA schedule (12h) for two weeks. Measures of motivation to self-administer oxycodone, hyperalgesia, analgesia and tolerance were performed for the whole duration of the experiment in >500 rats. These data are shown more in specific in a sister paper [23] from our laboratory, that aims to characterize addiction-like behaviors in heterogeneous stock rats. In the present study we tested the efficacy of buprenorphine, methadone, and naltrexone on the motivation for oxycodone intake in both males and females in a within latin-square design, and the behavioral tests associated with each group.

## Materials and Methods

### Animals

Heterogenous (HS) rats (*n* = 539 [277 males, 262 females]) were obtained from Wake Forest (Leah Solberg-Woods’ lab) at 3-4 weeks old. They were quarantined for two - three weeks and afterward were assigned IDs, weighed, and handled for 10-15 minutes over 3 days. Rats were housed two per cage with (Teklad 7090, Lab-Grade Sani-chios, Envigo, WI) bedding and maintained on a 12/12h light/dark cycle (lights off at 8:00 AM) in a temperature (20–22C) and humidity (45%–55%) controlled vivarium, with *ad libitum* access to (Teklad 8604, Lab Rodent Diet, Envigo, WI) food and tap water. All the procedures were performed two hours into the dark cycle, conducted in strict adherence to the National Institutes of Health Guide for the Care and Use of Laboratory Animals and were approved by the University of California, San Diego Institutional Animal Care and Use Committee.

### Drugs

Oxycodone (National Institute on Drug Abuse, Bethesda, MD) was dissolved in 0.9% sodium chloride (Hospita, Lake Forest, IL, USA). It was self-administered intravenously at a dose of 150 ug/0.1ml/kg. Buprenorphine hydrochloride (Sigma Aldrich) was administered at the dose 0.5 mg/kg, (±) – Methadone hydrochloride (Sigma Aldrich) at the dose 3mg/kg and naltrexone hydrochloride (Sigma Aldrich) was administered at the dose 3mg.kg interperitoneally (IP).

Buprenorphine, methadone, naltrexone, and saline were administered 30min prior test phase. The doses of buprenorphine, methadone and naltrexone were selected as the lowest effective dose, based on preliminary data performed in our aboratory on Wistar rats. All treatments were performed in a within subject design.

### Intravenous catheterization

The animals were anesthetized by inhalation of 3% isofluorane, and intravenous catheters were aseptically inserted into the right jugular vein using a modified version of a procedure that was described previously [24-26]. The catheters were flushed daily with heparinized saline (10 U/ml of heparin sodium; American Pharmaceutical Partners, Schaumburg, IL, USA) in 0.9% bacteriostatic sodium chloride that contained 20 mg/0.2 ml of the antibiotic Cefazolin (Hospira, Lake Forest, IL, USA).

### Oxycodone self-administration

Each self-administration session was initiated by the extension of two retractable levers into the operant chamber (29 cm x 24 cm x 19.5 cm; Med Associates, St. Albans, VT, USA). Self-administration sessions occurred in 2h sessions for 4 days (ShA), and 12-h daily sessions (LgA), starting at the beginning of the dark phase of the light/dark cycle, for 14 total sessions (5 sessions/week, Monday–Friday). Responses on the right active lever were reinforced on a fixed-ratio 1 (FR-1) schedule by intravenous oxycodone (150 μg/0.1 ml/infusion) administration that was infused over 6 s, followed by a 20-s timeout (TO20 s) period that was signaled by the illumination of a cue light above the active lever. Responses on the left inactive lever were recorded but had no scheduled consequences. The animals had access to food during the 12-h sessions. To test the motivation for oxycodone self-administration, rats were tested on a progressive ratio (PR) schedule of reinforcement at the end of each phase (once after ShA and once after LgA, see Fig 1 for detailed timeline), in which the response requirements for receiving a single reinforcement increased according to the following equation: [5e^(injection number x 0.2)^] − 5.

**Figure 1.**
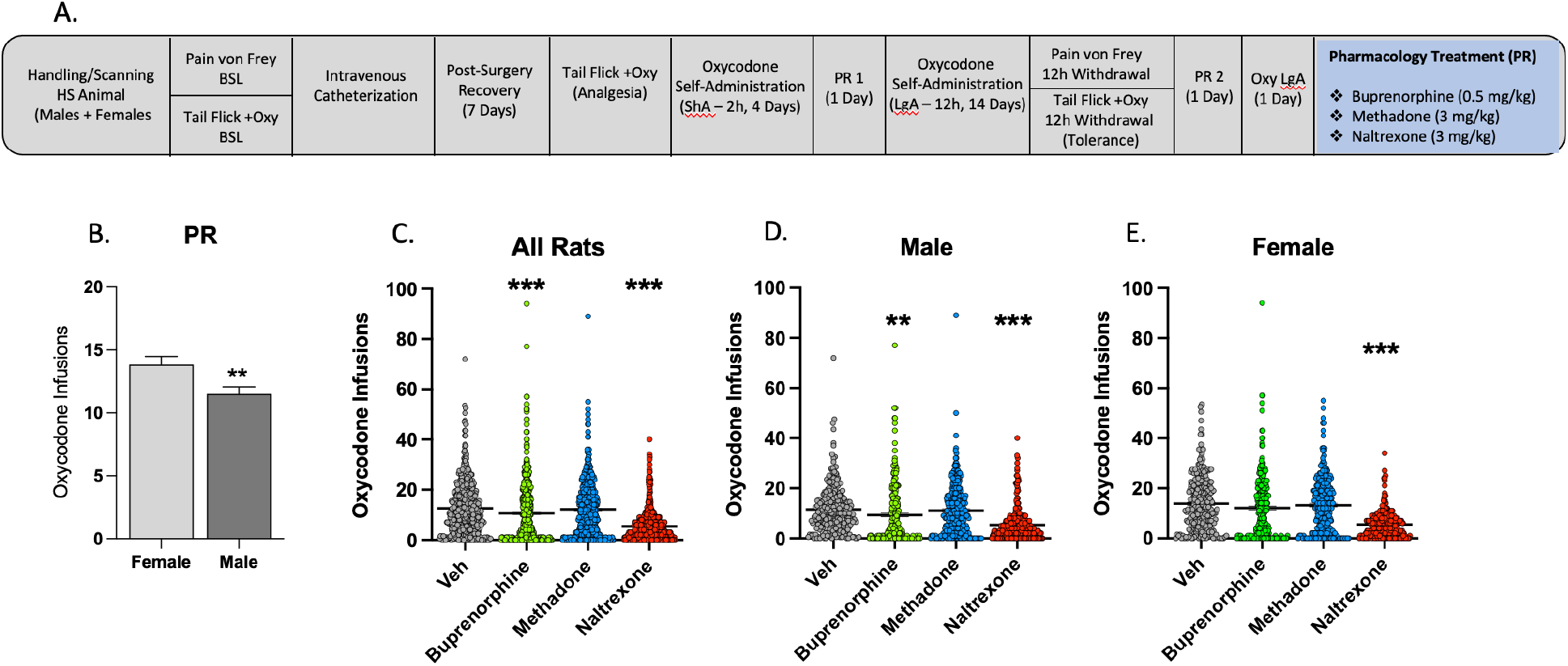
Effect of pharmacological treatment in a within latin-square design with buprenorphine, methadone and naltrexone on the motivation for oxycodone self-administration. A) Timeline of the behavioral paradigms. B) Oxycodone infusions during the last progressive ratio (prior treatment) in males and females. C) Effect of pharmacological treatment with buprenorphine, methadone and naltrexone in all rats. D) results show pharmacological treatment in males and E) in females. (** p < 0.01; ***p<0.001)

This resulted in the following progression of response requirements: 1, 1, 2, 2, 3, 3, 4, 4, 6, 6, 8, 8, 10, 11, 12, 13, 14, 15, 16, 17, etc +1 till 50, 60, 70, 80. 90, 100. The breakpoint was defined as the last ratio attained by the rat prior to a 60-min period during which a ratio was not completed, which ended the experiment. A block of fresh wood (3×3×3cm) was placed in the self-administration boxes during the oxycodone self-administration session.

Animals were intraperitoneally (IP) injected with either buprenorphine 0.5 mg/kg, methadone 3 mg/kg, naltrexone 3 mg/kg or saline 1ml/kg in a counterbalanced order, 30 min before the beginning of the PR session, to evaluate the effects of buprenorphine, methadone, and naltrexone on the motivation for oxycodone self-administration.

### Statistical Analysis

Statistics were calculated using GraphPad Prism 8.0. All values are expressed as mean ± SEM. The self-administration data were analyzed using repeated measures analysis of variance (ANOVA) of the number of infusions that were earned during the escalation interval. The effects of buprenorphine, methadone, and naltrexone on the escalation of oxycodone self-administration were analyzed using appropriate one-way ANOVAs (within-subjects).

Significant effects in the ANOVA were followed by the Dunnett post hoc test. Values of *p < 0.05* were considered statistically significant.

## Results

### 1 Effect of buprenorphine, methadone, and naltrexone on the motivation for oxycodone self-administration

To test the effect of buprenorphine (0.5mg/kg), methadone (3mg/kg) and naltrexone (3mg/kg) on the motivation for oxycodone self-administration, (n=539) rats were administered IP in a within latin-square design, 30 min prior the progressive ratio (PR) session. The number of oxycodone rewards and break points were recorded. Animals were left undisturbed in their home cages the day following treatment, to allow for the drugs to wash out. Statistical analysis one-way ANOVA showed an overall significant treatment effect (one-way ANOVA, *F*_3,538_ = 93.15, *p* <0.001; Fig. 1C). The Dunnett multiple-comparison test revealed a significant decrease of oxycodone rewards in the rats treated with buprenorphine (*p*<0.01) and with naltrexone (*p*<0.001) compared to the vehicle group. We then analyzed the efficacy of these treatments in both males and females. Statistical analysis one-way ANOVA showed an overall significant treatment effect (one-way ANOVA, *F*_3,276_ = 37.05, *p* <0.001) in males (Fig 1D). The Dunnett multiple-comparison test revealed a significant decrease of oxycodone rewards in the rats treated with buprenorphine (*p*<0.01) and with naltrexone (*p*<0.001) compared to the vehicle group. In the female group (Fig 1E), overall treatment was significant (one-way ANOVA, *F*_3,261_ =57.91, *p* <0.001). The Dunnett multiple-comparison test revealed a statistically significant decrease of oxycodone rewards only when the rats were treated with naltrexone (*p*<0.001) compared to the vehicle group.

To test whether there was any sex difference in the motivation for oxycodone self-administration, we analyzed the data of the last PR session, prior to the treatment administration. Student’s t-test analysis revealed a significant increase in oxycodone rewards during PR in females compared to males (*t*_537_ = 2.69, *p*<0.01) suggesting a contribution of biological sex to the motivation for oxycodone self-administration in rats (Fig 1B).

### 2 Effect of buprenorphine, methadone, and naltrexone on the motivation for oxycodone self-administration in the resilient and mild population

Using the addiction index [23], animals were ranked from low to high addiction-like behaviors. We differentiated between resilient, mild, moderate and severe addiction like behaviors in the whole population, by dividing them in 4 equal quartiles. We then analyzed the effect *of buprenorphine, methadone, and naltrexone on the motivation for oxycodone self-administration in the two lower quartiles that correspond to HS rats with lower addiction index (resilient and mild)* as shown in Figs 2A and 2E),. *We then evaluated the effect of pharmacological treatment in males and females in each quartile*.

**Figure 2.**
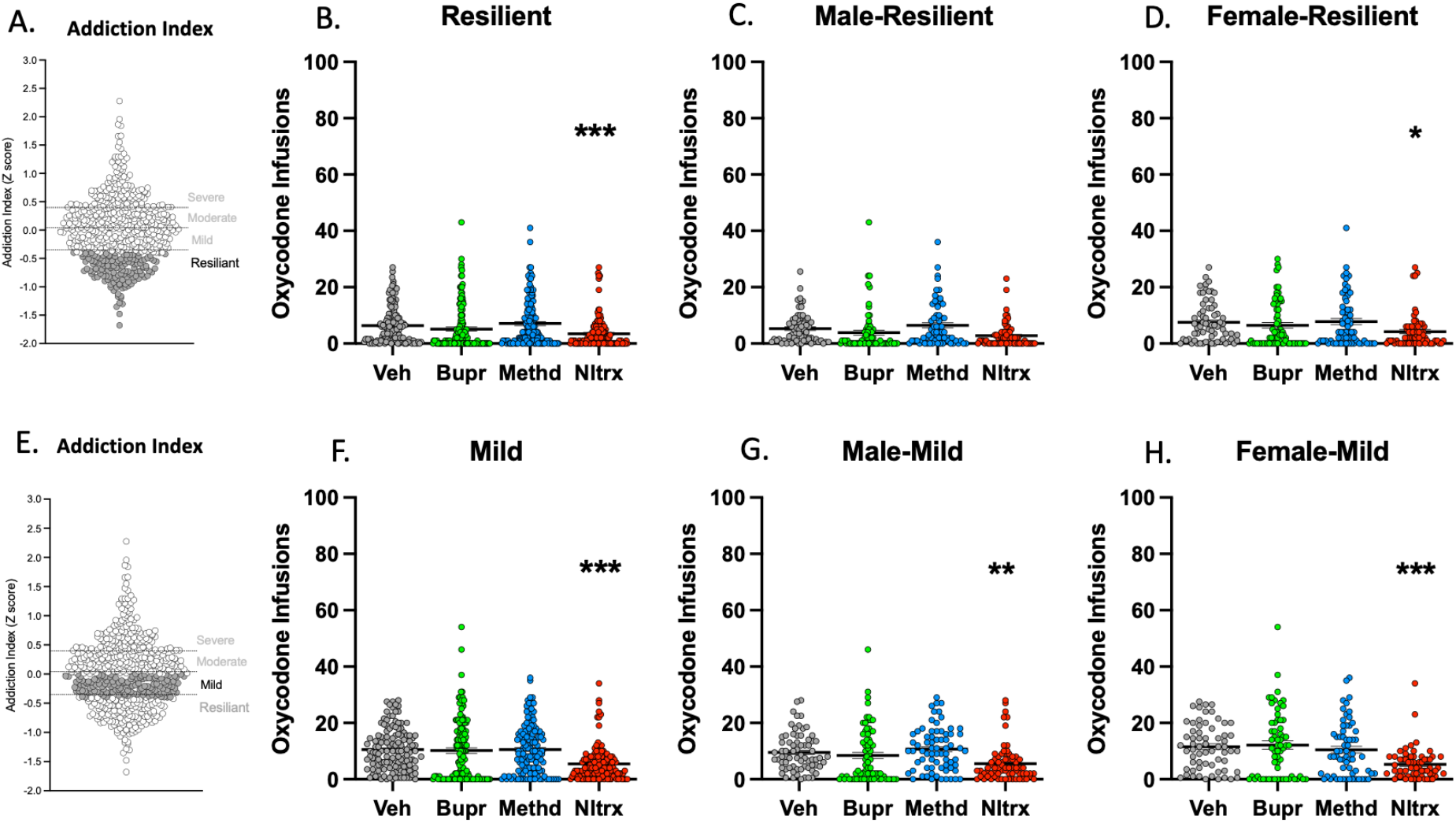
Effect of buprenorphine, methadone, and naltrexone on the motivation for oxycodone self-administration in the resilient and mild subpopulation. A) Representation of the resilient group (grey) based on the addiction index z-score. B) shows the pharmacological treatment with buprenorphine, methadone and naltrexone in the whole resilient group. C) shows results in males D) shows results in females. E) Representation of the mild group (grey) based on the addiction index z-score. F) Effect of pharmacological treatment with buprenorphine, methadone and naltrexone in the mild group. G) shows results in males H) shows results in females in the mild group (* p < 0.05;** p < 0.01; ***p<0.001).

Statistical analysis one-way ANOVA showed an overall significant treatment effect in the resilient group (*F*_3,135_ = 9.82, *p* <0.001) (Fig 2B). The Dunnett multiple-comparison test revealed a significant decrease of oxycodone rewards when rats were treated with naltrexone (*p*<0.001) compared to the vehicle group, but no effect of either buprenorphine or methadone was observed. We then analyzed the efficacy of these treatments in the resilient male and female subgroups. One-way ANOVA showed an overall significant treatment effect (*F*_3,67_ = 6.04, *p* <0.01) in male (Fig 2C), and in the female rats (*F*_3,67_ =4.42, *p* <0.01). Post hoc Dunnett’s test revealed a significant decrease of oxycodone rewards from naltrexone (*p*<0.05) compared to the vehicle group in females (Fig 2D) but none in males.

Overall one-way ANOVA showed significant treatment effect in the mild group (*F*_3,133_ = 17.27, *p* <0.001) (Fig 2F), and post hoc analysis revealed a significant decrease of oxycodone rewards after administration with naltrexone (*p*<0.001). We then analyzed the efficacy of these treatments in the mild males and females subgroups as shown in (Fig 2G and 2H), In both sexes, one-way ANOVA showed an overall significant treatment effect (*F*_3,70_ = 7.69, *p* <0.001) in males (Fig 2G) and (*F*_3,62_ =11.96, *p* <0.001) in females (Fig 2H). The Dunnett multiple-comparison test revealed a significant decrease of oxycodone rewards after treatment with naltrexone in both sexes (*p*<0.01) in male and (*p*<0.001) in females.

### 3 Effect of buprenorphine, methadone, and naltrexone on the motivation for oxycodone self-administration in the moderate and severe population

Here, we analyzed the effect *of buprenorphine, methadone, and naltrexone on the motivation for oxycodone self-administration in the two higher quartiles that correspond to HS rats with higher addiction index (moderate and resilient)* as indicated in Fig. 3A and Fig. 3E. *We then evaluated the effect of pharmacological treatment in males and females in each quartile*.

**Figure 3.**
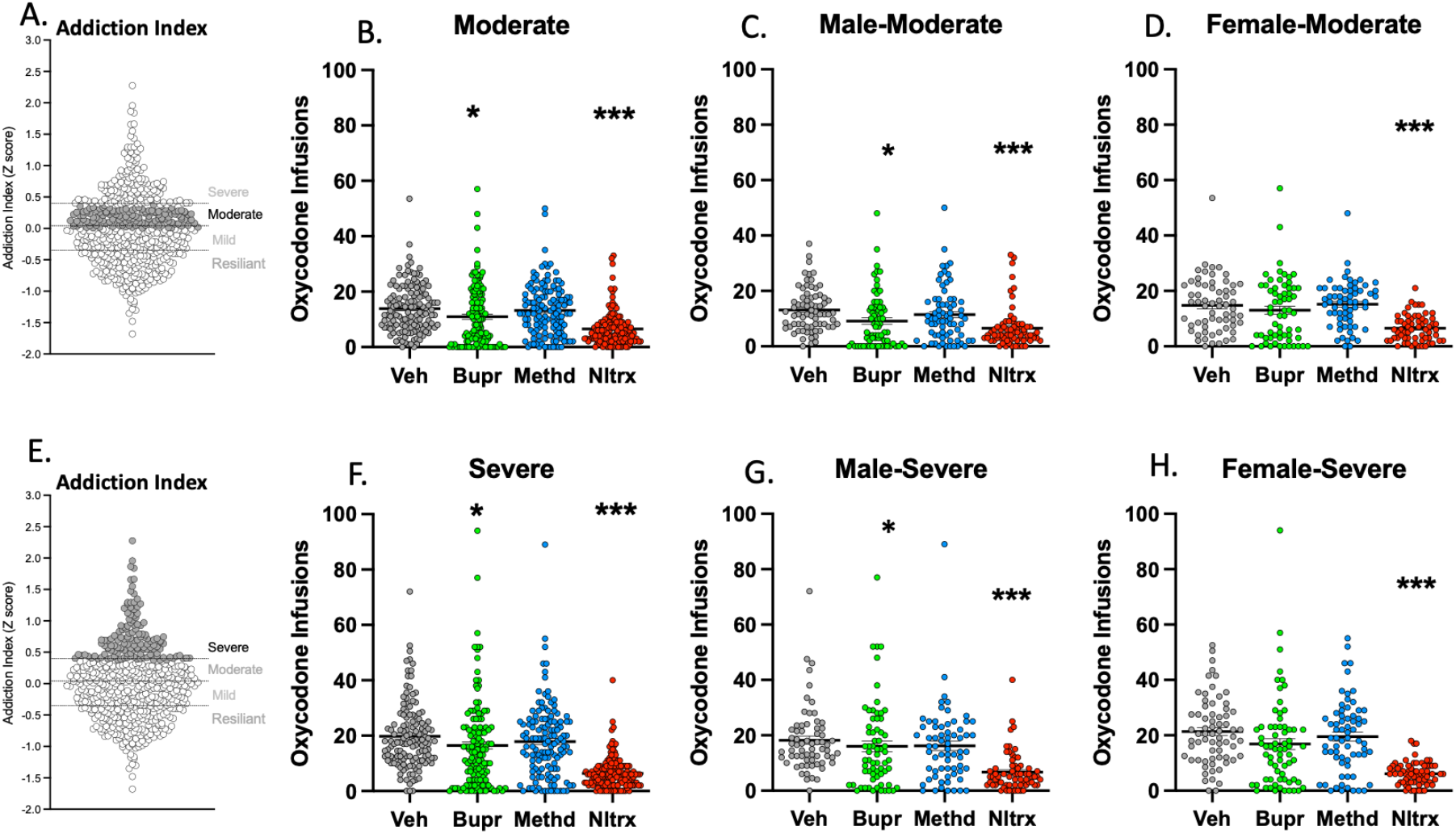
Effect of buprenorphine, methadone, and naltrexone on the motivation for oxycodone self-administration in the moderate and severe subpopulation. A) Representation of the moderate group (grey) based on the addiction index z-score. B) shows the pharmacological treatment with buprenorphine, methadone and naltrexone in the whole moderate group. C) shows results in males D) shows results in females. E) Representation of the severe group (grey) based on the addiction index z-score. F) Effect of pharmacological treatment with buprenorphine, methadone and naltrexone in the severe group. G) shows results in males H) shows results in females in the severe group. (* p < 0.05;***p<0.001)

**Figure 4.**
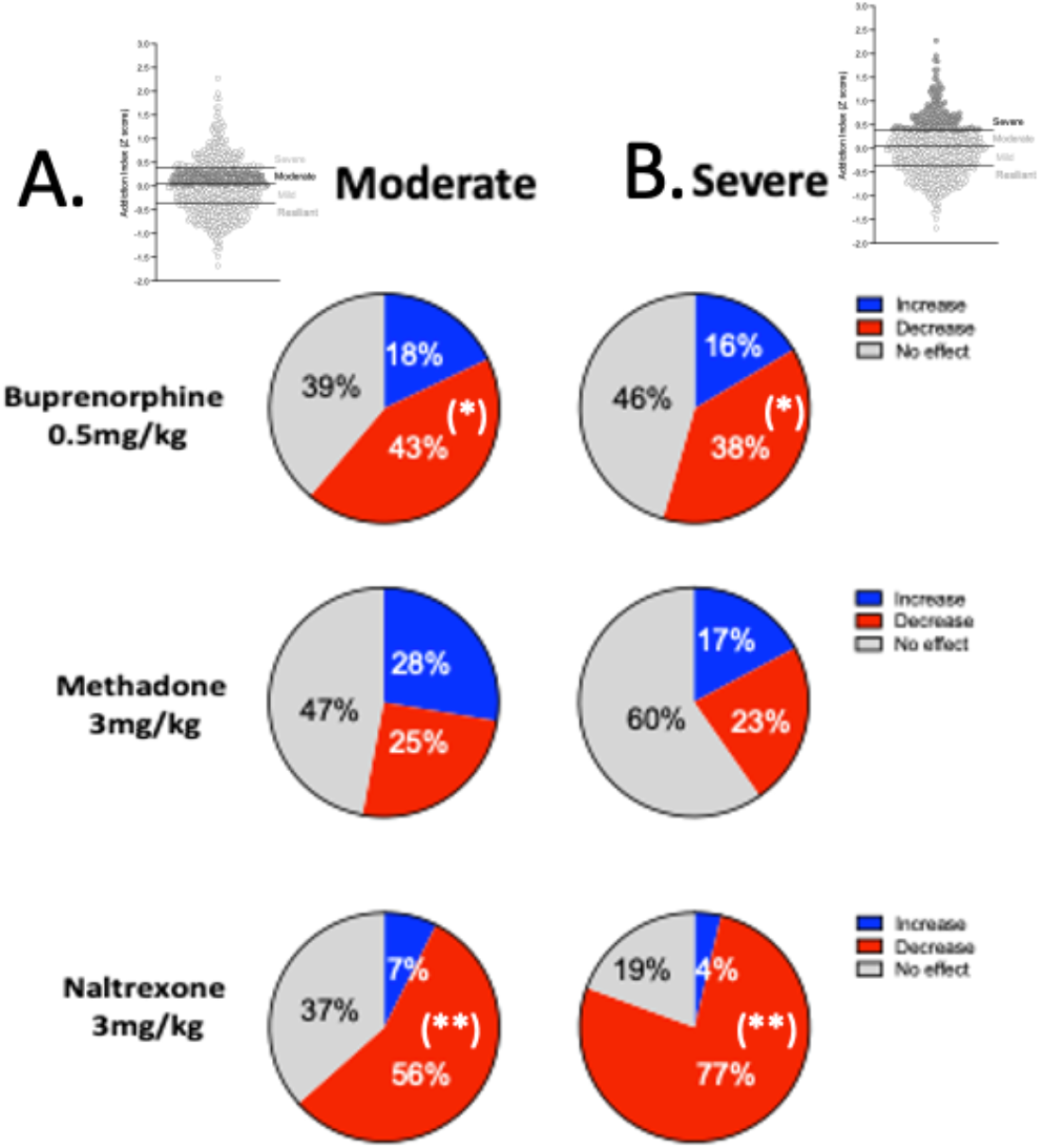
Pie chart of responders for each individual treatment in the A) moderate and B) severe groups. In grey is indicated the % of the population that did not respond to the treatment. In red is the % of the population that showed >50% decrease from the vehicle group and in blue the % of the population that showed increase responding of oxycodone self-administration in PR after treatment with buprenorphine, methadone or naltrexone (* p < 0.05;** p < 0.01; ***p<0.001)

Statistical analysis one-way ANOVA showed an overall significant treatment effect in the moderate group (*F*_3,134_ = 2.448, *p* <0.001) (Fig 3B). The Dunnett multiple-comparison test revealed a significant decrease of oxycodone rewards after treatment with naltrexone (*p*<0.001) and buprenorphine (*p*<0.05) compared to the vehicle group. We then analyzed the efficacy of these treatments in the resilient males and females of the moderate quartile. One-way ANOVA showed a significant treatment effect (*F*_3,71_ = 10.42, *p* <0.05) in males (Fig 3C) and in females (*F*_3,62_ =17.27, *p* <0.001) (Fig 3D). The Dunnett’s post hoc analysis revealed a significant decrease of oxycodone rewards after treated with naltrexone (*p*<0.001) and buprenorphine (*p*<0.05) in males, while statistically significant results were obtained only with the naltrexone treatment (*p*<0.001) in the female group of the moderate quartile.

Finally, one-way ANOVA showed an overall significant treatment effect in the severe group (*F*_3,133_ = 48.40, *p* <0.001) as shown in (Fig 3F). The Dunnett multiple-comparison test revealed a significant decrease of oxycodone rewards after treatment with naltrexone (*p*<0.001) and buprenorphine (*p*<0.05) compared to the vehicle group. We then analyzed the efficacy of these treatments in the severe male and female subpopulation. In both sexes, statistical analysis one-way ANOVA showed an overall significant treatment effect (*F*_3,65_ = 17.96, *p* <0.05) in males (Fig 3G) and (*F*_3,67_ =31.56, *p* <0.001) in females (Fig 3H). The Dunnett’s post hoc analysis revealed a significant decrease of oxycodone rewards after treated with naltrexone (*p*<0.001) and buprenorphine (*p*<0.05) in males, while statistically significant results were obtained only with the naltrexone treatment (*p*<0.001) in the female group of the severe quartile.

## Discussion

Opioid misuse remains a significant problem for our society. While currently available treatments (buprenorphine, methadone, and naltrexone) are effective in many patients, negative effects such as precipitated withdrawal, or respiratory depression from drug-drug interactions limit the effectiveness of these medications in the rest of the population. Thus, these is a need to find better treatments for opioid abuse and better translational models of preclinical research. Here we used HS rats, a unique genetically diverse population that self-administers oxycodone with high inter-individual and low intra-individual variability. This study provides results demonstrating the efficacy of naltrexone in reducing the motivation for oxycodone self-administration in the whole population of HS rats that escalated oxycodone intake. Acute treatment with naltrexone 3mg/kg significantly reduced the number of oxycodone rewards self-administered by both male and female rats. However, when we analyzed the data based on the addiction index of each individual animal, results showed that in the mild, moderate and in the severe addicted groups, this effect was maintained in male and female rats, but not in the resilient group. Naltrexone failed to significantly reduce the progressive ratio responses for oxycodone in males, while maintaining its effect in females. Naltrexone was efficacious in 56% of the population that classified as moderate addicted rats, and in 77% of the population in the severe addicted rats.

As expected, our data showed that buprenorphine (0.5 mg/kg) significantly reduced the motivation for oxycodone self-administration, but this effect was mainly attributed to males. In fact, when the entire population was divided into 4 groups based on their addiction index, buprenorphine maintained its efficacy only in the moderate (43%) and severe (38%) of the population, with mainly males’ response to buprenorphine’s effect.

We did not observe any statistically significant reduction in the motivation for oxycodone self-administration induced by methadone in any of the groups. Results showed that in the moderate and severe groups, there is a small percent of the population (25%) and (23%) respectively that reduced their progressive ratio responses after methadone administration.

The present study confirmed earlier findings [27] in which naltrexone reduced dose-dependently the responses for heroin in Rhesus monkeys and [28] in which naltrexone reduced the final ratios in heroin infusions in rats.

There have been relatively few studies that directly compare male and female addiction-like behaviors for oxycodone self-administration in preclinical models [5, 29-32]. In our study, the effect of naltrexone in reducing motivation for oxycodone self-administration was similarly reduced in male and female HS rats. A significant body of research has shown that buprenorphine and methadone reduce opioid use and improve retention to treatment [33, 34] While our data showed that buprenorphine reduced the progressive ratio of oxycodone self-administration, we also have evidence that this effect was maintained only in males and not in their female counterpart. One explanation to this observation may be that the higher escalation of oxycodone self-administration in females, that correlates to higher motivation for oxycodone intake in females compared to males, may have contributed to a desensitization of the μ-opioid receptor. Our data are in line with results from [19] showing that in males, buprenorphine increased the price sensitivity to fentanyl, by increasing the elasticity of demand for fentanyl.

These data suggest that buprenorphine may reduce demand by partially substituting for fentanyl or oxycodone at the μ-receptors, but also via actions at other receptor subtypes.

Moreover, buprenorphine significantly reduced motivation for oxycodone intake only in males in the moderate and severe group, suggesting that perhaps higher doses of buprenorphine would have been needed for the other less vulnerable subgroups or for the females.

Finally, ∼25% of individuals were sensitive to methadone, although methadone failed to reduce oxycodone self-administration during progressive ratio at the population level, suggesting that a higher dose may be required to observe population effects.

Altogether, the data shown in this study suggest that the efficacy of the medications used for the treatment of opioid addiction may vary between individuals with different opioid exposure, diverse addiction index and may be sex-specific. Further studies are needed to better understand the complicated pharmacology of these medications for the development of personalized medications for the treatment of opioid addiction.

### Limitations

One caveat of our study is that here, we did not perform a dose-effect of buprenorphine, methadone, and naltrexone, because we used the lowest efficacious dose that resulted from a pilot data performed in Wistar rats. Another caveat of this work is that these treatments were performed only in rats self-administering oxycodone. A control group of animals self-administering food or sucrose would have been necessary to test the behavioral specificity of these treatments. However, considering the large number of the population used for this study, a parallel experiment on the pharmacological effect of these medications on natural rewards does not justify the use of such a cohort of animals. Future studies need to be performed to address this issue in a different context of behavioral assays. We also did not evaluate any negative withdrawal symptoms that may have been induced by the treatments, specifically by naltrexone, because the time points of the experiments would have not been consistent due to the different individual responses during PR.

## Conclusion

This study provides preclinical evidence of therapeutic efficacy of naltrexone, buprenorphine, and methadone treatment on the reduction of the motivation for oxycodone intake. These pharmacological effects depend on the severity of the addiction state, while a large percentage of the population did not either respond to the treatment or increased their motivation for oxycodone intake. Further work is necessary to developing better drugs for the treatment of opioid addiction in different stages of its severity. This behavioral characterization was the first step in characterizing and understanding individual differences in a genetically diverse population. Future studies will combine these differences with the animals’ genetic and genomic data obtained through whole genome sequencing to perform a GWAS aimed to investigate gene variants that contribute to vulnerability and resilience to oxycodone addiction-like behavior (Chitre et al., 2020). This will then allow for the identification of specific genetic loci, estimation of heritability, investigation of genetic correlations and performance of phenome-wide association studies (PheWAS) and transcriptome-wide association studies (TWAS). All these studies will inform on individual differences in the sensitivity to FDA-approved. Finally, to facilitate investigating the biological origins of the difference to treatment samples of these behavioral and genetically characterized animals are collected and available free of charge through the oxycodone biobank [35].

## Acknowledgements

We thank the NIDA Center for GWAS in Outbred Rats and the Preclinical Addiction Research Consortium at UCSD.

